# Mutation induced disruption of domain coupling and interaction networks governs dysfunction of hDicer in recognizing pre-miRNA

**DOI:** 10.64898/2026.11.20.767749

**Authors:** Amrit Venkatesan, Atanu Maity, Prashasti Sinha, Ranjit Prasad Bahadur

## Abstract

Human Dicer (hDicer) is a multidomain RNase III enzyme central to RNA interference. Precise coordination between different domains in hDicer governs recognition and processing of double-stranded RNA substrate. While recent structural studies reveal distinct apo and dicing conformations, dynamics and energetics basis of hDicer function, particularly the effect of mutations in RNA recognition remains elusive. In this study, we investigate the mechanistic impact of two specific mutations in hDicer, A872T in platform domain and H1549Y in RIIIDa domain, in recognizing pre-miRNA. Our study shows that the wild-type RNA-bound complex exhibits a relatively confined conformational landscape, stable coordination between helicase and PAZ domains, and a persistent protein-RNA interaction network contributing to domain coupling. The mutant, A872T retains near-WT binding energetics but induces altered conformational sampling, domain coordination and long-range communication accompanied by selective redistribution of hDicer-pre-miRNA interaction networks. Conversely, H1549Y induces broader conformational heterogeneity, increased RNA exposure and redistribution of interdomain communication networks. Dynamic residue-network analysis identifies recurrent communication regions that connect mutation-associated perturbations with distal structural regions. This supports a mutation-specific mechanism, where the effect of local sequence alteration propagates through pre-existing intramolecular communication networks to alter domain coupling and RNA-interface organization without requiring global structural destabilization. A872T predominantly exhibits selective network rewiring with energetic compensation, whereas H1549Y produces broader communication and interaction-network redistribution accompanied by weaker energetic stabilization. Given the involvement of Dicer-dependent miRNA dysregulation in neurodegeneration and cancer, our findings provide a structural framework for understanding how domain-specific mutations may influence hDicer function and miRNA processing.

## 1. INTRODUCTION

Dicer is a central enzyme in the RNA interference (RNAi) pathway, responsible for processing precursor double-stranded RNA (dsRNA) into small regulatory RNAs including microRNAs (miRNAs) and small interfering RNAs (siRNAs) [1,2]. These small RNAs play essential roles in post-transcriptional gene regulation and are necessary for cellular homeostasis, development and disease progression [3]. Dicer is crucial for maintaining RNA-mediated gene regulation. Perturbation of its activity can have broad effects on cellular function. Human Dicer (hDicer) is a multidomain protein containing helicase domain, PAZ domain, platform domain, dual RNase III catalytic domain and double-stranded RNA-binding domain (dsRBD). All of these domains cooperate to recognize, position and cleave RNA substrates with high precision [2,4]. Recent high-resolution cryo-EM structures of hDicer reveals distinct conformations in apo (RNA-free) and RNA-bound dicing states, highlighting the importance of large-scale conformational rearrangements during substrate processing [1]. Apo hDicer adopts a relatively open and flexible architecture, whereas RNA binding promotes a more organized conformation that positions the substrate relative to the RNase III catalytic center [5]. In addition to these global rearrangements, local RNA features such as the GYM motif plays important role in selection of cleavage-site through coordinated interactions with dsRBD and RNase III domains [6]. Thus, hDicer function depends on the coordinated interplay of local interactions, domain coupling and long-range conformational dynamics.

Despite these structural insights, the dynamic and energetic mechanisms by which disease-associated mutations perturb hDicer function remain poorly understood. Mutations may preserve the overall fold while altering interaction networks, domain coupling or conformational landscapes, producing effects that may not be captured by static structural analysis or overall binding affinity alone [7]. Molecular dynamics (MD) simulations have provided insights into hDicer flexibility, domain motions and RNA recognition [8–11]. However, an integrated understanding on the effect of mutation-induced conformational changes with residue-level communication and protein-RNA interaction networks remains limited. The hDicer is significant for RNA processing, which is crucial for motor-neuron maintenance. Dysregulation of miRNA expression and processing has been reported in motor-neuron diseases, including Amyotrophic Lateral Sclerosis (ALS) [12–14]. This provides the rationale for analysing sequence and structural variations in hDicer and the effect of mutations in RNA processing.

In this study, we identify three non-synonymous hDicer variants Y1835C, H1549Y and A872T from RNA-Seq dataset of ALS patients. These variants are located within the RNase IIIDb (RIIIDb), RNase IIIDa (RIIIDa) and platform regions of hDicer, respectively. We further investigate the effects of H1549Y and A872T on pre-miRNA recognition using structural modelling and MD simulations. By integrating conformational analysis, collective motions, protein-RNA interactions, dynamic residue networks and binding energies, we find that these two mutations produce distinct perturbations rather than a common global destabilization of interaction networks between hDicer and pre-miRNA. A872T selectively alters domain coupling and long-range communication while retaining near-wild-type RNA-binding energetics. In contrast, H1549Y causes broader redistribution of protein-RNA interactions, increased RNA solvent exposure and weaker binding energetics. Dynamic residue-network analysis further identified recurrent communication regions, associating the mutation sites with distal regions of hDicer. This supports a mechanism in which local mutations propagate through intramolecular communication networks to alter RNA-interaction organization affecting RNA processing.

## 2. METHODS

### 2.1 Identification and selection of hDicer variants

RNA-Seq data from ALS samples (GSE124439) were analysed to identify sequence variation within hDicer [15–17]. The ALS dataset comprised 162 samples after exclusion of samples annotated as “Other Neurological Disorders” [18]. The sequencing reads were converted into FASTQ format using fasterq-dump [19], followed by quality assessment with FastQC [20]. Adapter sequences and low-quality bases were subsequently removed using Cutadapt [21]. The quality-filtered reads were then aligned to the human reference genome (GRCh38.p14) using STAR aligner v2.7.10 [22]. Following this, variant calling was performed using bcftools [23]. Variants were retained based on a variant quality score (QUAL > 30) and adequate read depth (DP > 20), followed by assessment of allele balance, base quality, strand representation and reproducibility across samples. The retained high-confidence variants were subjected to gene-based annotation, performed using ANNOVAR [24]. Within the *DICER1* gene, three heterozygous non-synonymous variants p.Tyr1835Cys (c.5504A>G), p.His1549Tyr (c.4645C>T) and p.Ala872Thr (c.2614G>A) were identified. H1549Y and A872T were selected for further structural and MD analysis as because their mutation sites were resolved in the available apo (PDB ID: 7XW3) and dicing-state (PDB ID: 7XW2) structures [25].

### 2.2 Preparation of wild-type and mutant structures

Cryo-EM structures of hDicer (PDB IDs: 7XW3 and 7XW2) were obtained from the Protein Data Bank [26]. Both the structures contain unresolved or missing regions, which were modelled using AlphaFold3 (DeepMind) [27]. The predicted fragments were structurally aligned and integrated into the experimental structures using PyMOL (version 2.5.0, Schrödinger, LLC) [28]. The resulting hybrid structures were carefully inspected to ensure proper backbone continuity and absence of steric clashes, followed by local geometric optimization. Point mutations A872T and H1549Y were introduced in the dicing-state structure using PyMOL. Appropriate side-chain rotamers were selected based on minimal steric hindrance and favourable local interactions. The resulting mutant structures were subsequently prepared for MD simulations.

### 2.3 System setup and production run

All the simulations were performed using GROMACS (version 2024.2) [29]. Protein and RNA parameters were assigned using an all-atom force field AMBER99SB-ILDN with compatible nucleic acid parameters [30]. Each system was placed in a triclinic simulation box with a minimum solute-box distance of 1.5 nm. The systems were solvated using SPC water model and the topology files were updated accordingly [31]. Sodium (Na^+^) and chloride (Cl^-^) ions were added to neutralize the system and achieve a physiological salt concentration of 0.15 M. Energy minimization was performed using the steepest-descent algorithm until the maximum force was below 1000 kJ/mol/nm. NVT equilibration was performed at 300 K using the V-rescale thermostat with heavy-atom positional restraints. NPT equilibration was performed at 1 bar using the Parrinello-Rahman barostat [32]. Equilibration phase of NVT and NPT was performed for 100 ps and 500 ps, respectively. The Apo-hDicer, WT-Complex, A872T and H1549Y systems were simulated for 1 μs each with two independent replicates per system. Hydrogen-bonds (H-bonds) were constrained using LINCS algorithm [33]. Long-range electrostatics were treated using Particle Mesh Ewald with a short-range cut-off distance of 10 Å for short-range interactions [34]. Periodic boundary conditions were applied in all directions. A timestep of 2 fs was used to save the trajectories.

### 2.4 Analysis of structural stability, compactness and flexibility

The overall structural stability of each system was evaluated using root mean square deviation (RMSD) of backbone atoms using gmx rms. RMSD profiles were used to assess equilibration behaviour and long-timescale conformational stability [35]. Structural compactness was quantified using the radius of gyration (Rg) computed with gmx gyrate. Local flexibility and dynamic behaviour were analysed using root mean square fluctuation (RMSF) using gmx rmsf. The values were computed on a per-residue basis to identify regions exhibiting enhanced flexibility or rigidity [36]. Time evolution of secondary structures was analysed using the DSSP algorithm implemented via gmx dssp [37,38].

### 2.5 Principal component analysis, free energy landscape and domain coordination

Large-scale collective motions were characterized using principal component analysis (PCA) [39]. The covariance matrix of atomic positional fluctuations was computed using gmx covar and the principal components were extracted through eigenvalue decomposition. Projection of trajectories along the dominant eigenvectors was performed using gmx anaeig, allowing identification of major conformational motions and comparison of dynamic behaviour across systems. The conformational space was further analysed by constructing free energy landscapes (FELs) based on projections along the first two principal components. Free energy surfaces were generated using gmx sham [29]. These landscapes were used to identify stable conformational basins and transitions between metastable states, providing insights into the energetic organization of the system [40,41]. PCA-derived collective motions were additionally visualized using porcupine plots. The dominant eigenvectors were mapped onto the corresponding protein structure to represent the direction and relative magnitude of the principal collective motions. The resulting vectors were used to compare the spatial distribution and directionality of large-scale motions among the Apo-hDicer, WT-Complex, A872T and H1549Y mutants. Interdomain rearrangements were quantified by measuring distances between key domains, specifically the helicase and the PAZ domains. Index groups corresponding to these domains were defined using gmx make_ndx and distances between their center of mass were calculated using gmx distance.

### 2.6 Dynamic residue network

Dynamic residue network analysis was performed to characterize residue-level communication in hDicer and identify pathways linking mutation sites to distal RNA-interacting regions [42–46]. Each residue was represented by its Cα atom, and residue pairs were connected when their Cα atoms were within 7.5 Å and showed dynamic correlation of |C| ≥ 0.60. Edges with ≥ 60% contact occupancy were retained for every tenth frame. Edge weights were assigned according to correlation strength; stronger correlations correspond to shorter network distances. Network properties, including degree of nodes, degree centrality, betweenness, closeness and eigenvector centrality were evaluated to assess the molecular interactions. Residues within high betweenness centrality were classified as potential communication intermediates. Weighted shortest paths between mutation sites and persistent protein-RNA interface residues were subsequently determined. Network visualizations were generated using Cytoscape v3.10.4 [47]. The network topology was used to characterize mutation-associated communication pathways between hDicer and pre-miRNA.

### 2.7 Hydrogen bond and solvent accessible surface area

Intermolecular H-bonds were analysed using gmx hbond. H-bonds were identified based on default geometric criteria including donor-acceptor distance and angular constraints [48]. The number of H-bonds was monitored throughout the simulation trajectories to evaluate interaction stability and binding integrity [49]. Solvent-accessible surface area (SASA) was calculated using gmx sasa to evaluate residue exposure and identify RNA-interacting regions [50,51]. Differences in SASA (ΔSASA) relative to WT-Complex were computed to assess mutation-induced perturbations at the interface.

### 2.8 Binding free energy and residue decomposition

Binding free energies between hDicer and pre-miRNA were estimated using the MM/PBSA approach implemented through the gmx_MMPBSA [52–54]. The total binding free energy (Δ) was decomposed into van der Waals (vdW), electrostatic, polar solvation and non-polar contributions. Additionally, residue-wise energy decomposition was performed to identify key interaction hotspots and quantify the impact of mutations on the energetic organization of the RNA-protein interface.

## 3. RESULTS

### 3.1 Identification and selection of hDicer variants

We have identified three non-synonymous substitutions within hDicer through variant calling analysis of RNA-Seq dataset obtained from 162 samples of ALS patients. These protein-altering variants include Y1835C, H1549Y and A872T, which are located at RIIIDb, RIIIDa and platform region of hDicer, respectively. For subsequent molecular characterization, H1549Y and A872T are selected due to their occurrence in the experimentally resolved regions of the available PDB structures of Apo-hDicer and WT-Complex. Two selected variants are therefore used to investigate mutation-specific effects on conformational dynamics, interdomain coordination and protein-RNA interactions. The domain organisation of hDicer comprises helicase, pincer-like, DUF283, D-linker, platform, PAZ, connector, C-linker, RIIIDa, RIIIDb and dsRBD (Fig. 1A). Helicase and DUF283 domains contribute to substrate recognition and RNA interactions. Platform and PAZ domains recognise and anchor the 5′-phosphate and the 3′-overhang ends of pre-miRNA, respectively. RIIIDa and RIIIDb domains form the catalytic centre responsible for cleavage of both the RNA strands, while dsRBD contributes to double-stranded RNA binding. The three-dimensional arrangement of these domains in Apo-hDicer and hDicer-pre-miRNA complex is shown in Fig. 1B and C, respectively. This highlights the structural organisation of hDicer and its interaction with the pre-miRNA substrate.

**Fig. 1.**
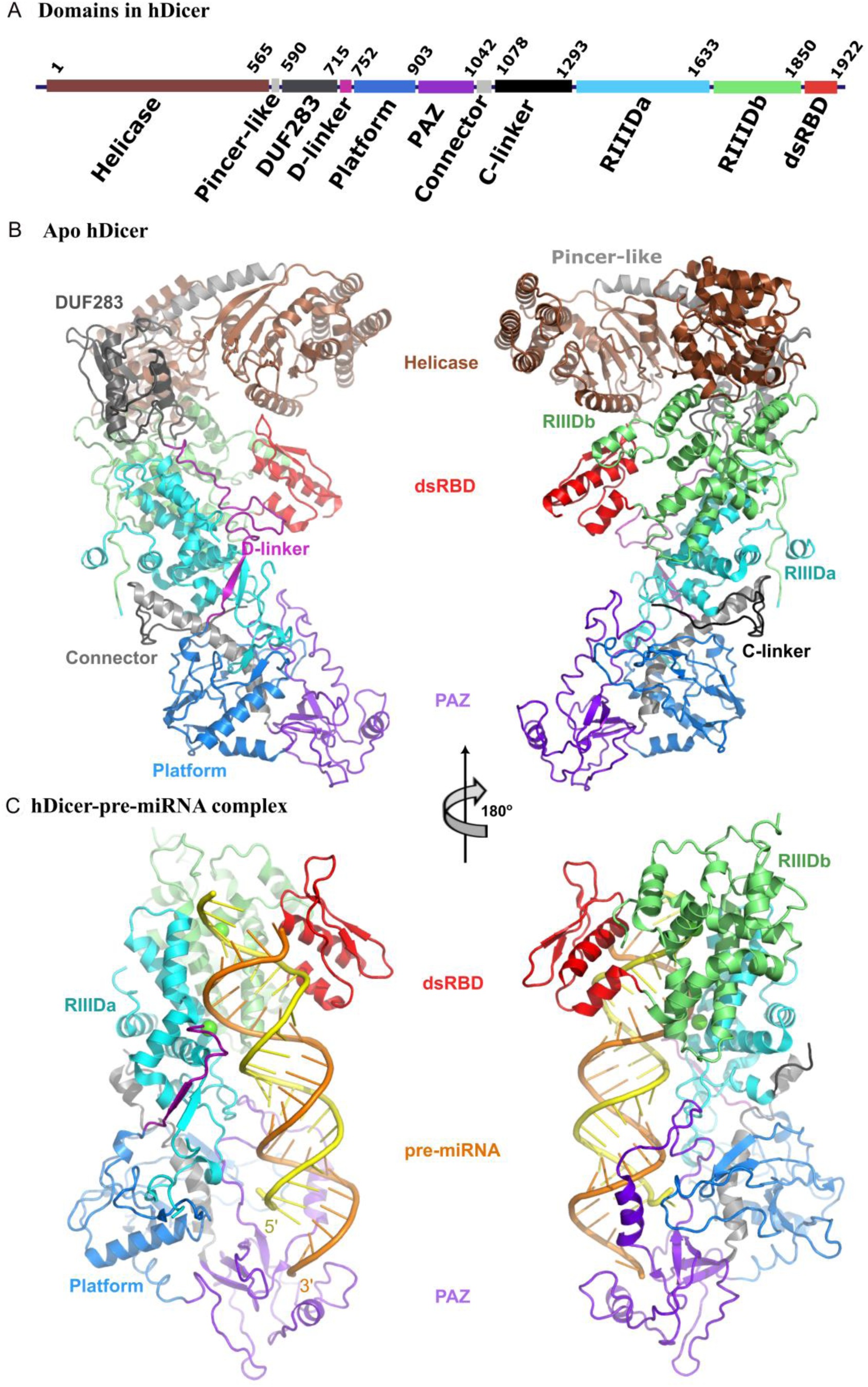
Distribution of different domains in hDicer (A). Three dimensional structure of Apo-hDicer (B) and hDicer-pre-miRNA complex (C).

### 3.2 Structural stability, compactness and residue-level dynamics

All systems show relatively stable RMSD profiles (Fig. 2A; Supplementary Table S1). WT-Complex exhibits the lowest RMSD (∼0.47 nm), indicating enhanced structural stability upon binding pre-miRNA (Supplementary Table S1). In contrast, Apo-hDicer shows higher RMSD (∼0.62 nm) with broader fluctuations, consistent with increased conformational flexibility in the absence of pre-miRNA. A872T shows RMSD comparable to that of WT-Complex (∼0.43 nm), suggesting minimal impact of mutation on the overall structural stability. In contrast, H1549Y shows slightly higher RMSD (∼0.52 nm) and pronounced fluctuations, indicating that mutation reduces stability and increases conformational heterogeneity [55].

**Fig. 2:**
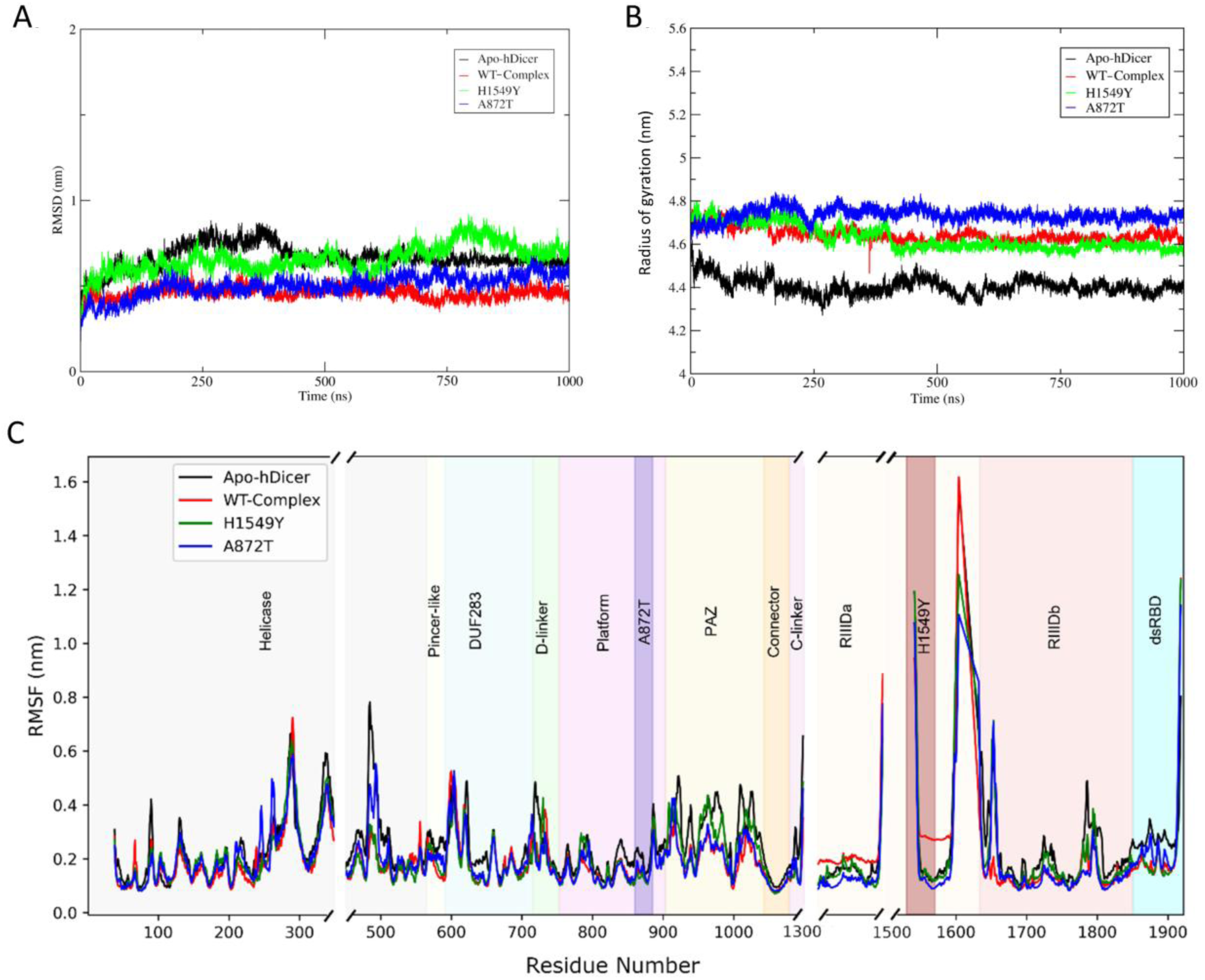
Structural stability, compactness and residue-level flexibility of Apo-hDicer, WT-Complex and mutant systems. (A) Backbone root mean square deviation, (B) Radius of gyration and (C) Residue-wise root mean square fluctuation. Different domains of hDicer are highlighed using different colour schemes. Discontinuous residue streches in hDicer are presented with gaps.

Rg analysis reveals differences in the overall compactness of the systems (Fig. 2B; Supplementary Table S1). Apo-hDicer exhibits the lowest Rg (∼4.45 nm), indicating a relatively compact conformation. On the other hand, WT-Complex shows an increased Rg (∼4.66 nm) upon binding pre-miRNA, reflecting structural expansion associated with substrate accommodation. Mutant A872T displays the highest Rg (∼4.72 nm) with relatively stable fluctuations, indicating relatively expanded conformation compared to WT-Complex. However, mutant H1549Y maintains an Rg (∼4.65 nm), close to that of WT-Complex.

RMSF profiles reveal domain-specific differences in residue flexibility across the systems (Fig. 2C). The helicase domain shows moderate fluctuations with localised peaks around residues 280 to 320. Apo-hDicer exhibits relatively higher fluctuations around residues 490 to 500. On the other hand, WT-Complex and H1549Y show reduced fluctuations in this region. Pincer-like, DUF283 and D-linker regions display broadly comparable flexibility. Platform domain, including the A872T mutation site, also shows similar RMSF profiles across the systems. Moderate fluctuations are observed in PAZ domain with slightly higher values in Apo-hDicer and H1549Y. A prominent fluctuation peak is observed in RIIIDa region around residues 1350 to 1450, particularly in WT-Complex. In the same region, WT-Complex shows a pronounced peak (1.6 nm) around residues 1600 to 1650, while Apo-hDicer displays intermediate behaviour and two mutants show reduced peaks. Compared to other systems, a slightly higher fluctuation is observed in dsRBD region of Apo-hDicer.

Stability of secondary structures is further assessed using DSSP (Supplementary Fig. S1A-D). The overall secondary structure remains conserved across all the systems throughout the simulation. WT-Complex maintains stable α-helices and β-sheets with minimal temporal variation. Apo-hDicer shows slightly increased variability in the loop and the turn regions, while both the mutants exhibit minor local variations in secondary structures. Our results do not show any major global secondary structure transitions in all the systems.

### 3.3 Conformational landscape, collective motions and interdomain coordination

The conformational landscapes of Apo-hDicer, WT-Complex, A872T and H1549Y are investigated using PCA with FEL constructed from projections along the first two principal components (Fig. 3). Apo-hDicer exhibits a broad conformational distribution with multiple clusters, indicating extensive sampling of different conformational states (Fig. 3 A(i)). In contrast, WT-Complex displays a relatively confined and continuous low-energy basin, suggesting a more organised conformational landscape following RNA binding (Fig. 3 B(i)). The mutant A872T exhibits two distinct low-energy basins, indicating the sampling of alternate conformational states compared to the continuous landscape of WT-Complex (Fig. 3 C(i)). The other mutant H1549Y shows a broader and more dispersed distribution with multiple low-energy basins extending across the conformational space (Fig. 3 D(i)). This pattern indicates increased conformational heterogeneity in H1549Y relative to WT-Complex and A872T. The corresponding PC1-derived porcupine plots reveal differences in the direction and spatial distribution of collective motions across the systems. Apo-hDicer and WT-Complex show distinct patterns of directional movements distributed across multiple structural regions (Fig. 3 A(ii) and B(ii)). WT-Complex exhibits comparatively coherent collective motions across major structural domains. A872T retains the overall framework of collective movements but shows altered orientations, particularly in linker and peripheral regions (Fig. 3 C(ii)). In contrast, H1549Y displays more heterogeneous directional movements across several structural regions, consistent with its broader conformational landscape (Fig. 3 D(ii)).

**Fig. 3:**
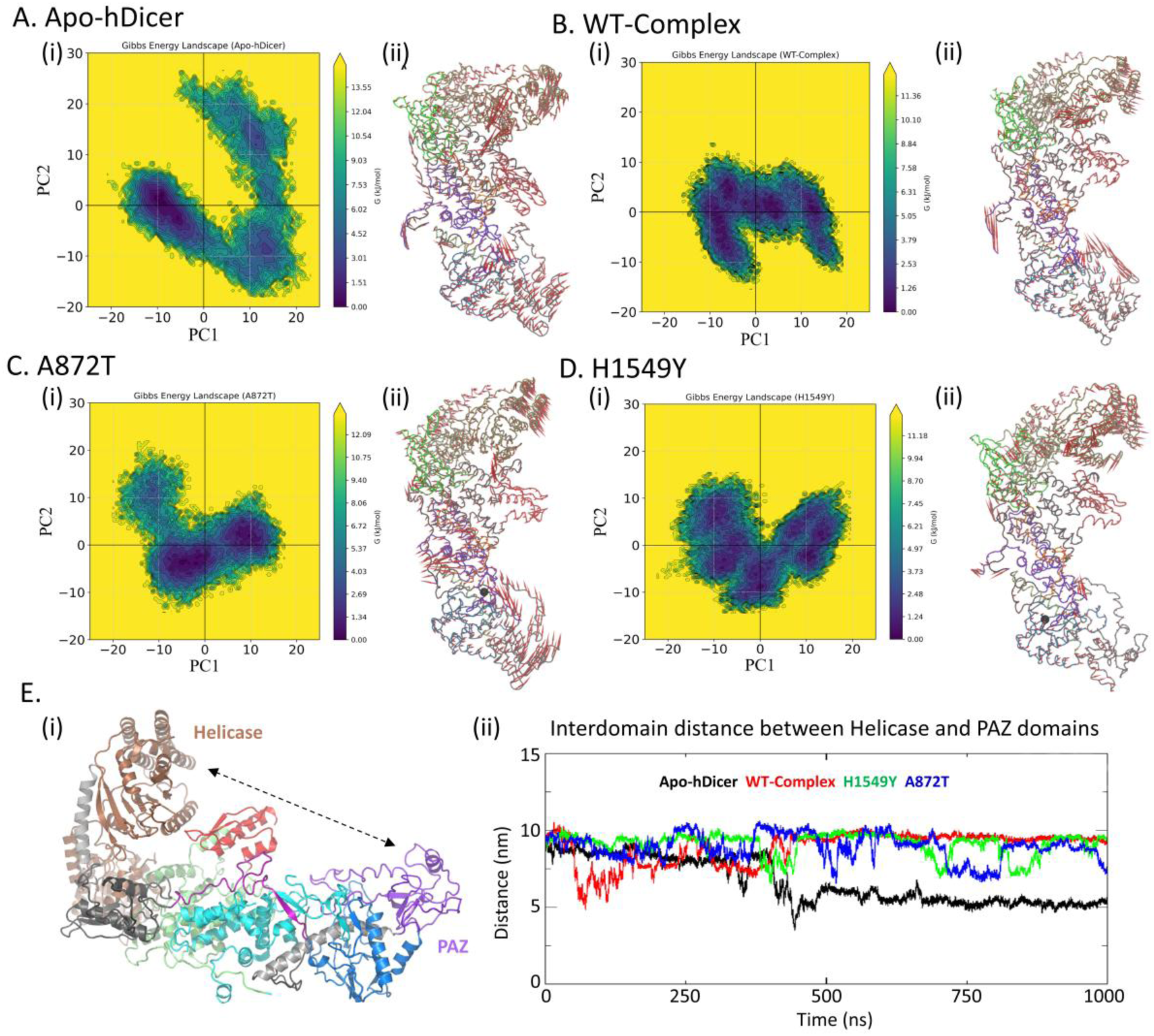
Conformational landscapes, collective motions and interdomain coordination of hDicer in (A) Apo-hDicer, (B) WT-Complex, (C) A872T and (D) H1549Y. (E) Interdomain-distance between helicase and PAZ domains.

Interdomain coordination between helicase and PAZ domains is evaluated by monitoring the distance between their center of mass (COM) over the simulation trajectory (Fig. 3 E(i)). WT-Complex maintains relatively stable interdomain distance, predominantly within 10 nm after the initial equilibrium phase with only minor fluctuations observed over time. Apo-hDicer exhibits a progressive decrease in interdomain distance, reaching approximately 6 nm after 400 ns (Fig. 3 E(ii)). This behaviour indicates a distinct domain arrangement compared to the RNA-bound system. Mutant A872T shows increased variability in interdomain distance, fluctuating between 7 nm and 10 nm. Multiple transitions between different distance states are observed throughout the trajectory. Similarly, H1549Y displays pronounced fluctuations in interdomain distance ranging from 7 nm to 10 nm with frequent transitions between conformational states.

### 3.4 Mutation-induced reorganisation of intramolecular communication networks

Dynamic residue-network (DRN) analysis is performed to investigate the effects of A872T and H1549Y on intramolecular communication pathways in hDicer. Communication networks starting from the points of mutation are shown in Figure 4 (A-F). A comparison of the network connectivity reveals that both mutations reorganised the existing communication pathway rather than causing a complete loss of network connectivity. Residues form interconnected networks spanning across the central and C-terminal regions in Apo-hDicer and WT-Complex with a different connectivity pattern. This indicates that binding pre-miRNA altered the communication landscape of Apo-hDicer. The initial phase of the communication pathway involving A872 with the RNA-interacting residues is similar in Apo-hDicer and WT-Complex. These include residues S874, A875, C877, F802 and I804 (Fig. 4A-B; Fig. 5A, B), predominantly located in platform domain. In contrast, the initial communication pathway in A872T includes residues Y1385, L1381, N1379, I1373 and D1308, located in RIIIDa domain (Fig. 4C). In the later phase of the communication, Apo-hDicer shows a considerable diversion connecting residues V720, P714, L712, H633 and D244 spanning helicase, linker and DUF283 regions (Fig. 4A and 5A). Conversely, WT-Complex and A872T follow a similar path connecting residues P1853, S1852, S1902, A1900, R1879 and T1873, located in dsRBD (Fig. 4B and 5B). Moreover, A872T retains connections with pre-miRNA-interacting residues I521, A518, I517 and Q501 in helicase domain. In addition, A872T establishes additional connections with R506, T519, V504 and L516 in helicase domain, which are absent in WT-Complex. The connection with F802 in platform domain is lost in A872T. These observations indicate selective modifications of the communication pathway to pre-miRNA-interacting regions.

**Fig. 4.**
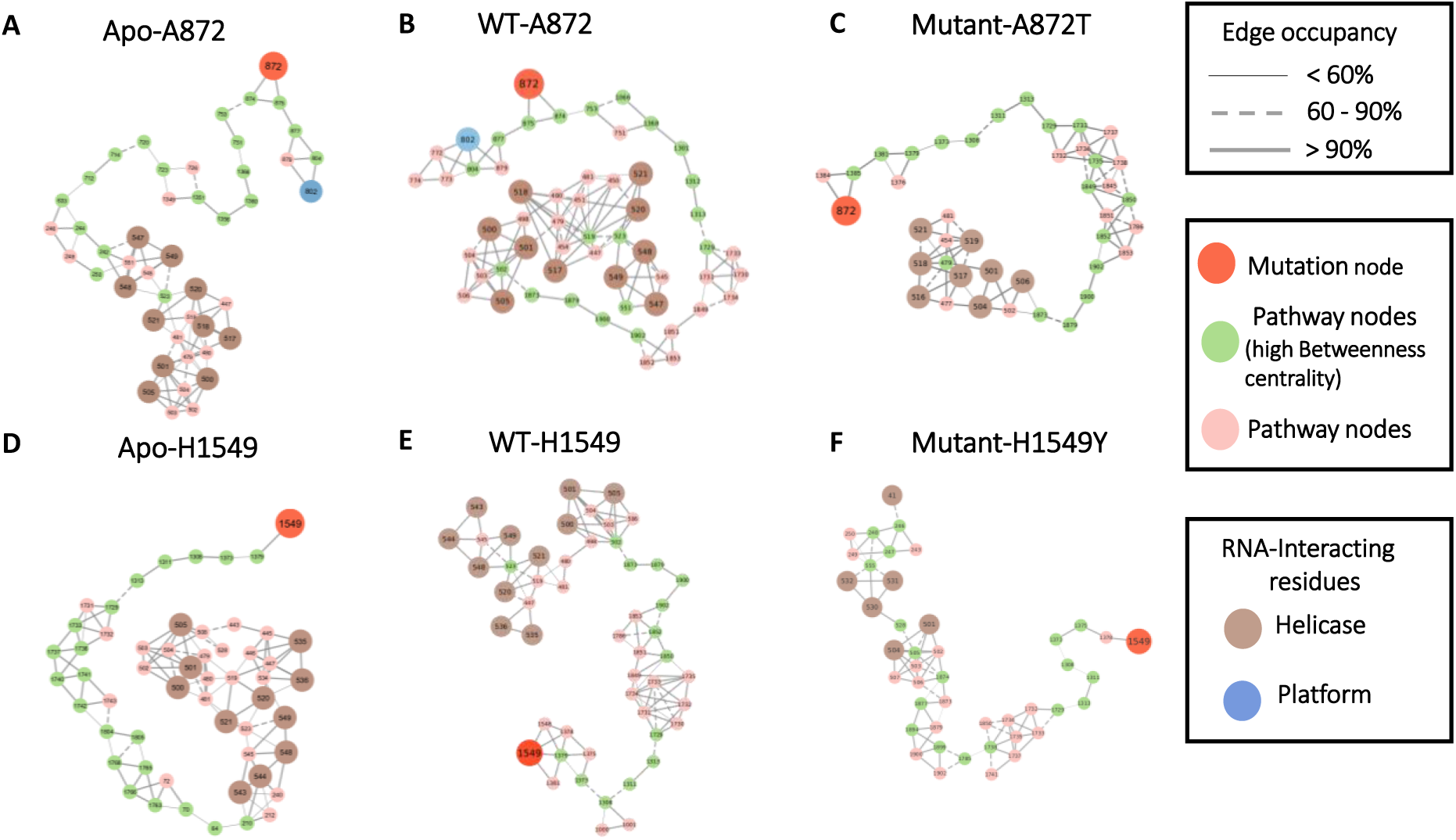
Dynamic residue network mapping of mutation-associated communication pathways in hDicer (A) Apo-A872, (B) WT-A872, (C) Mutant A872T, (D) WT-H1549, (E) Apo-H1549 and (F) Mutant H1549Y.

**Fig. 5.**
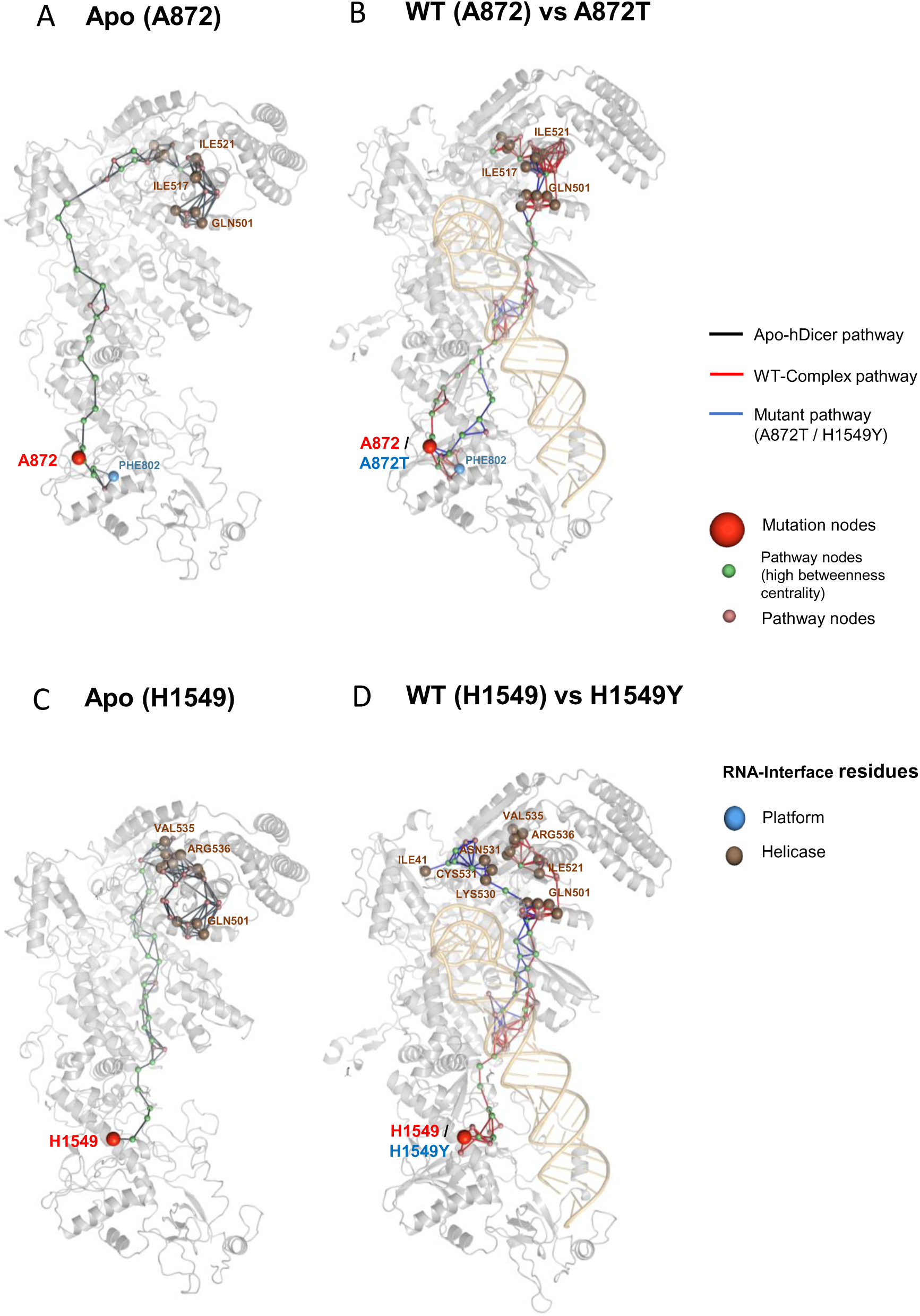
Three-dimensional representation of allosteric communication pathway from the site of mutation to the RNA-interacting residues for (A) Apo (A872), (B) WT (A872) vs A872T, (C) Apo (H1549) and (D) WT (H1549) vs H1549Y.

In case of H1549, pathway from the site of mutation in Apo-hDicer, WT-Complex and mutant H1549Y includes the residues D1375, I1373, D1308, N1311 and E1313, present in RIIIDa domain (Fig. 4D-F, Fig. 5C-D). This pathway extends towards residues P1729 to T1743 in RIIIDb. Compared to WT-Complex, H1549Y retains connection only with the pre-miRNA interacting residue Q501. Additionally, it establishes connection with a distinct cluster involving K530, C531, N532 and I41 in helicase region. This pattern indicates redistribution of communication towards alternate structural regions. Residues E1313 (RIIIDa), P1729 (RIIIDb), T1733 (RIIIDb), V1849 (dsRBD) and R1851 (dsRBD) repeatedly occur in WT and mutant networks. This suggests their potential role as communication intermediates linking structurally distant regions of hDicer. Therefore, A872T largely retains the interconnected communication framework of WT-Complex with selective changes in pathway connections. Whereas, H1549Y shows broader redistribution across distant residue clusters.

### 3.5 Protein-RNA interactions and interface solvent accessibility

Stability and organization of hDicer-pre-miRNA interface are investigated through H-bond analysis. The distribution of persistent hDicer-pre-miRNA H-bond occupancies reveals distinct interaction patterns across the complexes (Fig. 6 A, B). WT-Complex contains several H-bonds with higher occupancy, particularly involving residues in PAZ and dsRBD. Interactions such as TYR936-U73 and TYR976-U73 show high occupancy (∼ 82%), indicating persistent RNA anchoring. A872T retains several WT-like interactions along with selective redistribution of some H-bonds. Increased H-bond occupancies are observed for ARG1855-C19 and ARG996-C64, whereas the occupancy for SER995-G2 is reduced. H1549Y displays a broader redistribution of H-bond interactions, including increased persistence of interactions involving LYS975-U73 and ARG788-U65. These changes indicate that both mutations modify the distribution of persistent interactions between hDicer and pre-miRNA with a more dispersed interaction pattern in H1549Y.

**Fig. 6:**
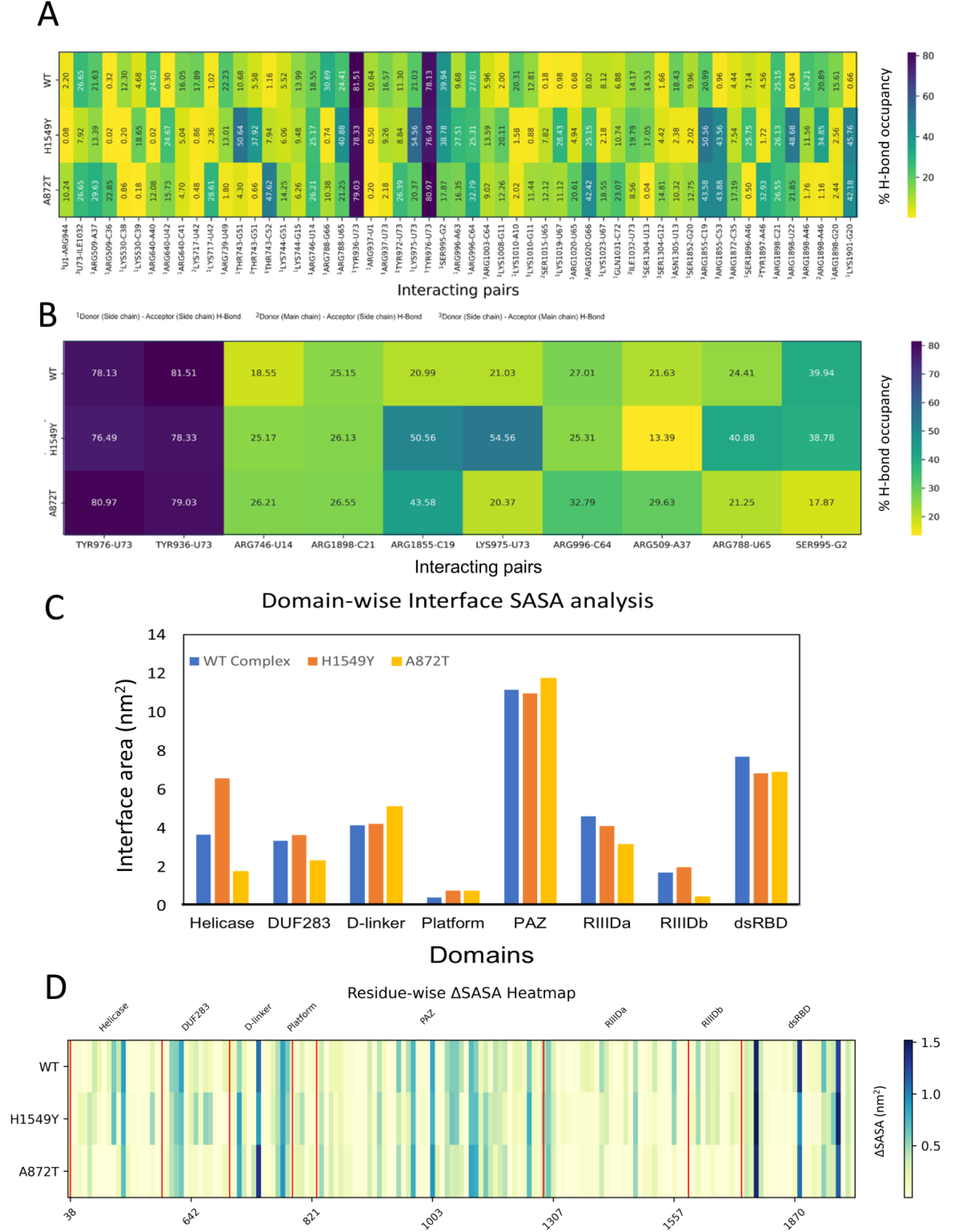
Mutation-dependent changes in protein–RNA H-bond interactions and interface solvent accessibility. (A) Percentage occupancy of intermolecular H-bonds between hDicer and pre-miRNA across WT-Complex, A872T and H1549Y, (B) Comparative heatmap of selected protein-RNA H-bond interactions, (C) Domain-wise contributions to the protein-RNA interface SASA in WT-Complex, A872T and H1549Y and (D) Residue-wise heatmap of change in interface area.

Domain-wise interface SASA further reveals differences in the contribution of individual domains to pre-miRNA binding (Fig. 6C; Table 1). In WT-Complex, PAZ domain contributes the largest interface area (11.16 nm^2^), followed by dsRBD (7.71 nm^2^) and RIIIDa (4.63 nm^2^). Compared to WT-Complex, the contribution of helicase domain increases from 3.67 nm^2^ to 6.58 nm^2^, while the contribution of RIIIDa decreases from 4.63 nm^2^ to 4.13 nm^2^ in H1549Y. In contrast, a reduced contribution by helicase (1.76 nm²) and RIIIDb (0.46 nm²) domains to the interface area is observed in A872T. Residue-level interface analysis also reveals mutation-dependent changes in solvent exposure at the hDicer-pre-miRNA interface (Fig. 6D; Supplementary Table S2). Prominent contributions are observed from R1855 and R1898 in dsRBD, and R996 and R1003 in PAZ. This indicates that the mutations alter the spatial organization of the hDicer-pre-miRNA interface rather than producing a uniform change across all interacting surface.

**Table 1.** Domain-wise protein-RNA interface SASA contributions. Contributions of each domain to hDicer-pre-miRNA interface area.

|  | Interface area (nm<sup>2</sup>) |  |  |
|---|---|---|---|
| Domain | WT<br>Complex | H1549Y | A872T |
| Helicase | 3.67 | 6.58 | 1.76 |
| DUF283 | 3.35 | 3.65 | 2.32 |
| D-linker | 4.16 | 4.24 | 5.12 |
| Platform | 0.43 | 0.79 | 0.75 |
| PAZ | 11.16 | 10.98 | 11.74 |
| RIIIDa | 4.63 | 4.13 | 3.17 |
| RIIIDb | 1.71 | 1.99 | 0.46 |
| dsRBD | 7.71 | 6.85 | 6.89 |
| **Total** | **36.82** | **39.21** | **32.21** |

### 3.6 Binding energetics and interaction network

The WT-Complex exhibits Δ of approximately -228.96 ± 33.58 kJ/mol with dominant contributions arising from electrostatic (-4248.21 ± 389.08 kJ/mol) and vdW (-322.42 ± 20.17 kJ/mol) interactions, partially offset by polar solvation energy (4376.40 ± 384.35 kJ/mol) (Fig. 7A-C; Table 2). In H1549Y, Δ increases to -197.43 ± 29.54 kJ/mol, indicating weaker binding relative to WT-Complex. Here, although contribution of vdW remains comparable (-330.76 ± 27.96 kJ/mol), contribution of electrostatic is less favourable (-4105.65 ± 383.52 kJ/mol) accompanied by an increased polar solvation (4275.57 ± 378.86 kJ/mol). Mutant A872T shows a Δ of -222.47 ± 26.18 kJ/mol, comparable to that of WT-Complex. In addition, A872T exhibits the most favourable electrostatic contribution (-4976.60 ± 491.53 kJ/mol), which is counterbalanced by a significantly higher polar solvation (5100.39 ± 472.95 kJ/mol), resulting in an overall Δ similar to that of WT-Complex. Residue-wise decomposition reveals heterogeneous energetic contributions across both protein and RNA components with mutation-dependent redistribution of favourable and unfavourable contributions (Fig. 7B, C). Specific nucleotides show strong favourable interactions, while others contribute unfavourably, indicating a non-uniform interaction landscape. The distribution of energetically significant nucleotides differs between WT-Complex and mutant systems, reflecting altered interaction patterns. Thus, the MM/PBSA analysis demonstrates that while electrostatic interactions dominate binding across all systems, the balance between electrostatic stabilization and reduced polar solvation determines the net binding affinity. Variations in residue-level contributions further indicate redistribution of interaction networks in the mutant systems.

**Fig. 7.**
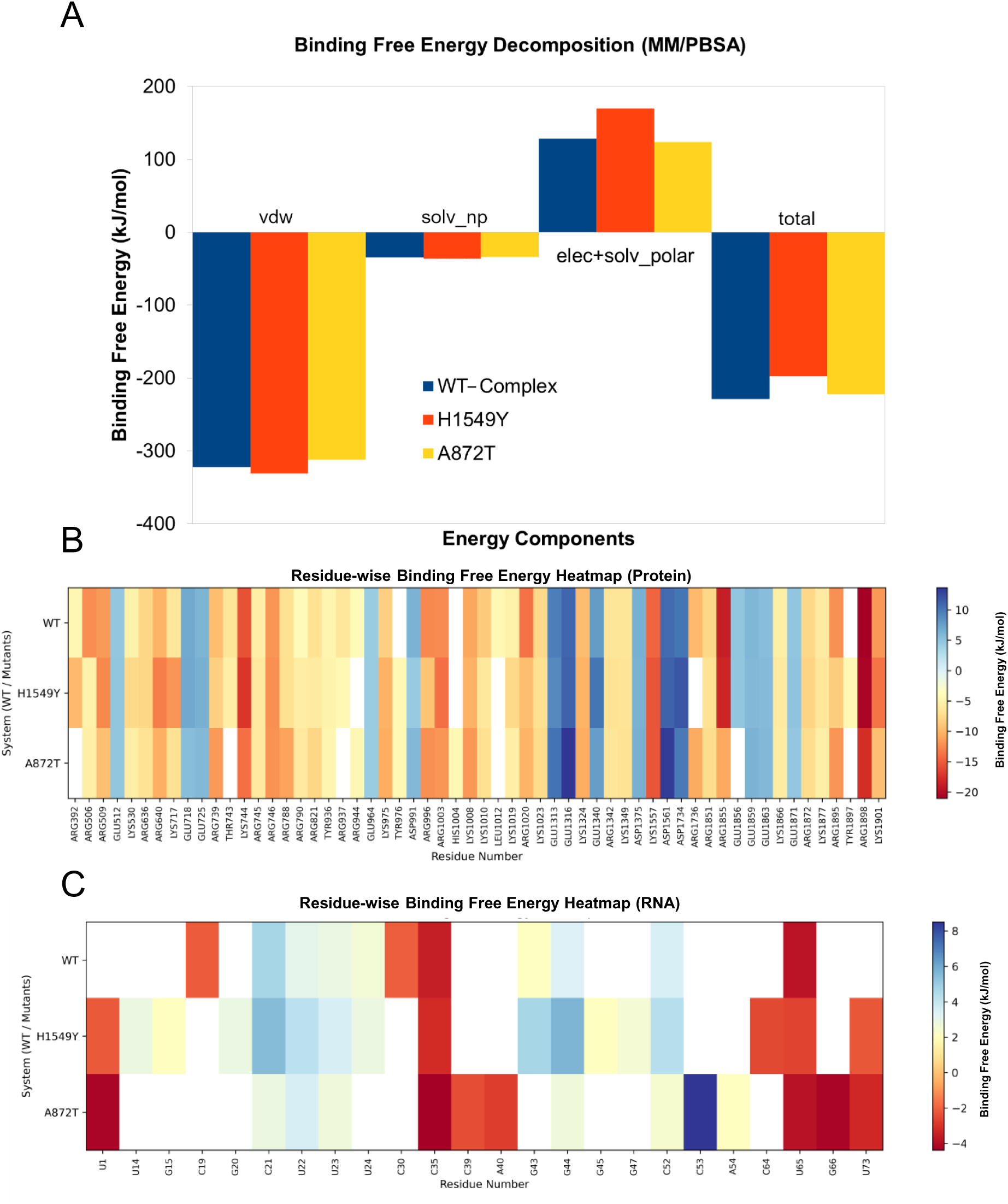
(A) Contribution of different energy terms in overall binding free energy. Decomposition of binding free energy of residues in hDicer (B) and nucleotides in pre-miRNA (C).

**Table 2.** Binding free energy components from MM/PBSA analysis.

| System | Elec (kJ/mol) | Vdw (kJ/mol) | solv_pol (kJ/mol) | solv_np (kJ/mol) | elec+solv_polar (kJ/mol) | Total ΔG (kJ/mol) |
|---|---|---|---|---|---|---|
| WT_Complex | -4248.21 ± 389.08 | -322.42 ± 20.17 | 4376.40 ± 384.35 | -34.73 ± 1.93 | 128.19 ± 4.73 | -228.96 ± 33.58 |
| H1549Y | -4105.65 ± 383.52 | -330.76 ± 27.96 | 4275.57 ± 378.86 | -36.60 ± 2.56 | 169.92 ± 4.66 | -197.43 ± 29.54 |
| A872T | -4976.60 ± 491.53 | -312.15 ± 16.66 | 5100.39 ± 472.95 | -34.11 ± 1.13 | 123.79 ± 18.58 | -222.47 ± 26.18 |

## 4. DISCUSSION

Dicer is a central component of miRNA biogenesis, essential for post-transcriptional gene regulation and the maintenance of neuronal homeostasis. miRNAs regulate several processes relevant to neuronal function, including synaptic activity, axonal maintenance and cellular stress responses. The importance of Dicer in motor neuron development and survival is supported by experimental studies [56,57]. Moreover, impaired Dicer-mediated miRNA processing and reduced miRNA abundance have been reported in ALS [58]. These findings provide a biological rationale for investigating whether sequence variations could influence structure and function of hDicer, contributing to altered RNA regulation in neurodegeneration. Although *DICER1* variants have been reported in genetic databases and disease-associated studies, their functional consequences remain poorly characterised. In the present study, we analyze two variants A872T and H1549Y of hDicer, which were identified from the RNA-Seq dataset of ALS patients. Both H1549Y and A872T are reported in the ClinVar database [59]. Although, A872T is classified as benign [60], H1549Y has not yet been functionally characterized and its clinical significance remains undetermined. However, the specific structural and functional consequences of these substitutions in hDicer remain poorly characterized. These variants are particularly relevant because of their occurrence in two distinct domains of hDicer. A872T occurs in platform domain and H1549Y occurs in RIIIDa, one of the catalytic domains. Location of these mutations provides an opportunity to examine how substitutions in distinct structural domains may affect domain coordination, intramolecular communication and RNA interactions. Therefore, the present study provides an initial computational assessment of their potential impact on hDicer structure, protein-RNA interactions and intramolecular communication networks.

### 4.1 Mutation-dependent changes in hDicer dynamics and RNA recognition

Comparison of Apo-hDicer and WT-Complex highlights the importance of substrate-associated conformational organisation. WT-Complex exhibits a relatively confined conformational landscape and stable helicase-PAZ coordination compared to Apo-hDicer. This is consistent with the cryo-EM study of hDicer-pre-miRNA, which demonstrates RNA-induced rearrangements of helicase, PAZ and dsRBD domains to position the substrate within the catalytic pocket [25]. A872T, located in platform domain, alters conformational sampling while maintaining overall structural stability and several WT-Complex-like RNA interactions. However, selective redistribution of interface contacts, including ARG1855-C19 and ARG996-C64, suggests changes in substrate organisation. The MM/PBSA binding affinity of A872T (-222.47 ± 26.18 kJ/mol) is comparable to that of WT-Complex (-228.96 ± 33.58 kJ/mol), indicating largely preserved energetic stabilisation. This suggests that the mutation-induced conformational changes do not affect the overall stability. In contrast, H1549Y induces broader conformational and interaction-network rearrangements along with weaker energetic stabilisation (-197.43 ± 29.54 kJ/mol) in hDicer. In H1549Y, the occupancies of H-bond interactions LYS975-U73 and ARG788-U65 increase relative to WT-Complex. Since H1549Y is located in RIIIDa, these changes may affect the organisation of hDicer-pre-miRNA complex. Previous studies have shown that interactions involving the platform, PAZ and dsRBD are important for pre-miRNA recognition and cleavage-site selection [1,2]. The alteration in interdomain coordination upon H1549Y mutation can also impact the coordination between these domains, resulting in weaker protein-RNA interactions.

### 4.2 Intramolecular communication networks and a proposed mechanism of hDicer perturbation

Collective motion and DRN provide insights how mutations at distinct positions influence the functional organisation of hDicer. Both the mutations reorganise pre-existing communication pathways rather than disrupting overall network connectivity. The DRN analysis further highlights E1313 in RIIIDa, P1729 and T1733 in RIIIDb, and V1849 and R1851 in the dsRBD as potential communication intermediates linking structurally distant regions of hDicer. Mutation-dependent redistribution of intramolecular communication pathways in WT-Complex, A872T and H1549Y is illustrated in Fig. 8. A872T largely retains the interconnected network framework of WT-Complex but selectively alters communication routes and connections with RNA-interacting residues in the helicase domain. This is consistent with the relatively localised changes observed in its collective-motion analyses. In contrast, H1549Y produces a broader redistribution of communication pathways across distant residue clusters, consistent with its increased conformational heterogeneity, dispersed protein-RNA interactions and weaker estimated binding energetics.

**Fig. 8.**
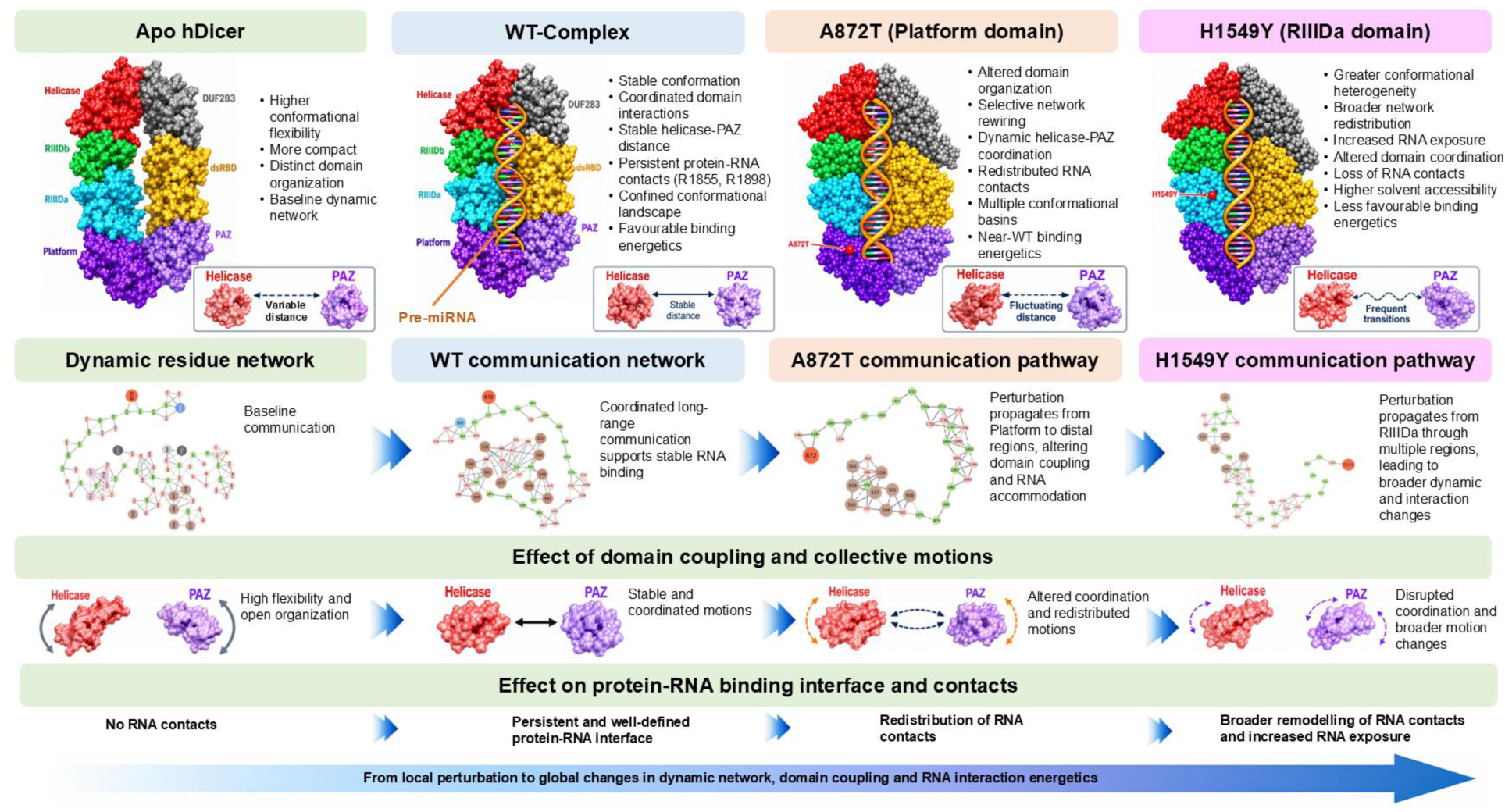
Integrated mechanistic framework.

These observations support the central hypothesis of the present study: mutations in distinct functional regions of hDicer can propagate through pre-existing intramolecular communication networks to modify interdomain coordination and the organisation of the protein-RNA interface without necessarily causing global structural destabilisation. A872T primarily induces selective network rewiring with energetic compensation, whereas H1549Y causes broader redistribution of communication and interaction networks accompanied by weaker energetic stabilisation. Such alterations may affect substrate positioning and conformational transitions required for efficient miRNA processing, potentially influencing downstream miRNA-regulated gene expression.

### 4.3 Effect of mutations in hDicer on miRNA biogenesis associated with neurodegeneration and cancer

Dicer-dependent miRNA biogenesis is essential for cellular homeostasis, including neuronal maintenance. Dysregulation of miRNAs has been implicated in neurodegenerative disorders, including ALS, where alterations in DICER-associated protein interactions have been reported [58,61,62]. In addition, cancer-associated mutations in the RIIIDa and RIIIDb of hDicer have been linked to impaired 5p-miRNA processing and altered gene expression. In particular, mutations in RIIIDb have been reported in pleuropulmonary blastoma and non-epithelial ovarian cancers [63–65]. The analyses of TCGA small RNA datasets indicate that RIIIDa mutations can also impair 5p-miRNA biogenesis [66]. These findings highlight the functional importance of catalytic domains of hDicer, suggesting that mutations in different structural regions may influence miRNA processing through distinct mechanisms. In this context, our findings reveal mutation-specific changes in hDicer interdomain coordination, intramolecular communication and protein-RNA interactions associated with A872T in the platform domain and H1549Y in RIIIDa. Although these structural and dynamic alterations provide a plausible mechanistic basis for altered hDicer function, their effects on miRNA processing and downstream gene regulation remain to be experimentally established.

## 5. CONCLUSION

This study demonstrates that A872T in platform domain and H1549Y in RIIIDa domain perturb hDicer-pre-miRNA interactions through distinct changes in its interdomain coupling, communication networks and binding energetics. The WT-Complex maintains a coordinated conformational state, stable domain organization and a persistent protein-RNA interaction network. Mutant A872T primarily induces selective reorganization of domain coordination and long-range communication while retaining near-WT binding energetics. This suggests that functional perturbation may arise from altered conformational organization and RNA accommodation rather than the loss of binding affinity. In contrast, mutant H1549Y produces broader redistribution of protein-RNA contacts and collective motions, increased RNA exposure, greater conformational heterogeneity and weaker energetic stabilisation of the pre-miRNA-bound complex. Dynamic residue-network analysis identifies recurrent regions, particularly E1313, P1729, T1733, V1849 and R1851, as communication intermediates linking mutation-associated perturbations to distal structural regions. Therefore, the present study supports a mechanistic model in which the effect of local mutations propagates through pre-existing intramolecular communication networks to alter domain coupling and RNA-interface organization without global structural disruption. The distinct effects of A872T and H1549Y further indicate that the mutations reshape the nature and extent of hDicer perturbation with a potential consequence of pre-miRNA recognition. Alterations in hDicer interactions and pre-miRNA recognition have been reported in neurodegeneration and cancer. Our findings provide structural and dynamic insights on how the specific mutations in hDicer may influence pre-miRNA recognition and RNA-interface organization. These observations provide a basis for further investigation into the functional relevance of hDicer mutants in neurodegeneration and cancer.

## Acknowledgment

AV and PS thank IIT Kharagpur for their fellowship and infrastructure support. AM and RPB thank IIT Kharagpur for infrastructure support. Both AM and RPB thank DBT, India for National Network Project (Grant no. BT/PR40182/BTIS/137/88/2025).

## Author contributions

RPB and AV conceptualized the study. AV, AM and RPB designed the study. AV, AM and PS performed the work, data analysis and wrote the initial draft of the manuscript. All authors read and approved the final manuscript. RPB analyzed data, reviewed and edited the manuscript, overall supervised the project, and arranged funding and resources.

## Supplementary information

Available as Supplementary Fig. S1 and Supplementary Table S1-S2.

## Ethics approval

Not applicable

