## Supplementary material for "Mutation induced disruption of domain coupling and interaction networks governs dysfunction of hDicer in recognizing pre-miRNA"

for

Ph: +91-3222-283790

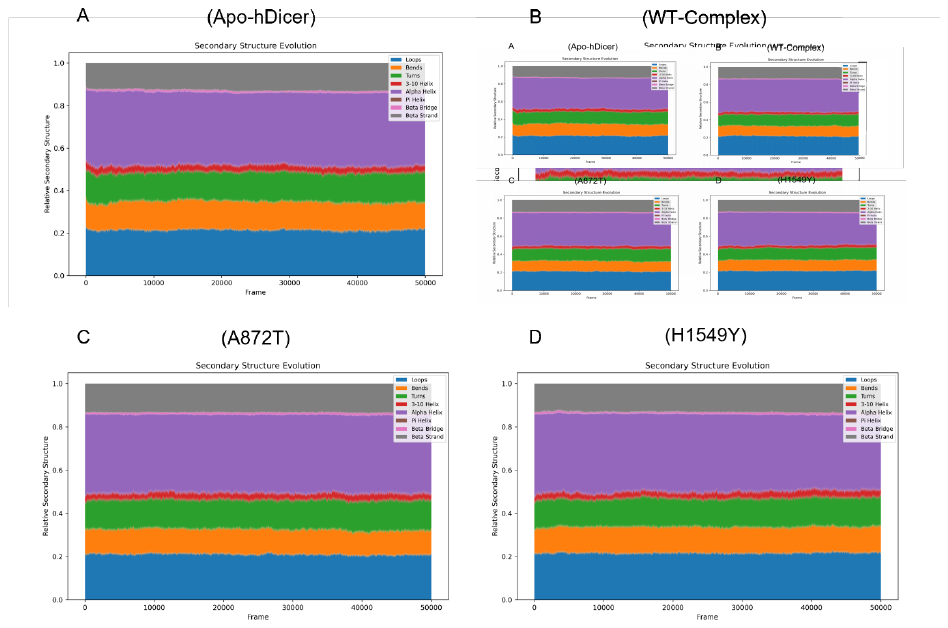

Fig. S1. Secondary structure stability analysis (A) Apo, (B) wild-type, (C) A872T mutant and (D) H1549Y mutant.

Table S1: Mean and standard deviation of the replicate systems.

|  |  | Rg (nm) | RMSD (nm) | RMSF (nm) |
| --- | --- | --- | --- | --- |
| Apo | R1 | 4.41 ± 0.038 | 0.67 ± 0.069 | 0.25 ± 0.151 |
|  | R2 | 4.51 ± 0.031 | 0.57 ± 0.041 | 0.22 v 0.137 |
| WT | R1 | 4.64 ± 0.032 | 0.46 ± 0.067 | 0.20 ± 0.133 |
|  | R2 | 4.70 ± 0.027 | 0.48 ± 0.067 | 0.21 ± 0.149 |
| H1549Y | R1 | 4.63 ± 0.058 | 0.66 ± 0.080 | 0.20 ± 0.134 |
|  | R2 | 4.69 ± 0.025 | 0.39 ± 0.059 | 0.21 ± 0.122 |
| A872T | R1 | 4.74 ± 0.029 | 0.50 ± 0.063 | 0.20 ± 0.131 |
|  | R2 | 4.68 ± 0.019 | 0.36 ± 0.034 | 0.19 ± 0.130 |

*R1: Replicate 1; R2: Replicate 2

Table S2. Residue-wise protein-RNA interface ΔSASA.

| H1549 | | | A872T | | | Complex_WT | | |
| --- | --- | --- | --- | --- | --- | --- | --- | --- |
| Domain | Residue number | ∆SASA (nm^2^) | Domain | Residue number | ∆SASA  (nm^2^) | Domain | Residue number | ∆SASA  (nm^2^) |
| Helicase | 38 | 0.028 | Helicase | 38 | 0.054 | Helicase | 38 | 0.045 |
|  | 39 | 0.003 |  | 39 | 0.037 |  | 39 | 0.018 |
|  | 43 | 0.181 |  | 41 | 0.002 |  | 43 | 0.039 |
|  | 44 | 0.143 |  | 43 | 0.111 |  | 243 | 0.001 |
|  | 45 | 0.004 |  | 44 | 0.011 |  | 246 | 0.077 |
|  | 46 | 0.088 |  | 46 | 0.017 |  | 247 | 0.001 |
|  | 128 | 0.001 |  | 243 | 0.011 |  | 390 | 0.02 |
|  | 131 | 0.03 |  | 245 | 0.041 |  | 391 | 0.02 |
|  | 132 | 0.381 |  | 246 | 0.343 |  | 392 | 0.128 |
|  | 133 | 0.015 |  | 395 | 0.001 |  | 393 | 0.132 |
|  | 134 | 0.373 |  | 397 | 0.002 |  | 394 | 0.214 |
|  | 135 | 0.099 |  | 430 | 0.001 |  | 395 | 0.351 |
|  | 136 | 0.109 |  | 431 | 0.001 |  | 396 | 0.038 |
|  | 137 | 0.101 |  | 437 | 0.001 |  | 397 | 0.168 |
|  | 140 | 0.002 |  | 438 | 0.001 |  | 430 | 0.001 |
|  | 157 | 0.003 |  | 506 | 0.179 |  | 431 | 0.001 |
|  | 160 | 0.085 |  | 507 | 0.005 |  | 434 | 0.005 |
|  | 161 | 0.221 |  | 509 | 0.683 |  | 436 | 0.005 |
|  | 162 | 0.005 |  | 510 | 0.146 |  | 437 | 0.218 |
|  | 163 | 0.146 |  | 511 | 0.012 |  | 438 | 0.588 |
|  | 242 | 0.079 |  | 512 | 0.016 |  | 439 | 0.001 |
|  | 243 | 0.154 |  | 527 | 0.001 |  | 440 | 0.001 |
|  | 246 | 0.382 |  | 529 | 0.01 |  | 441 | 0.001 |
|  | 247 | 0.076 |  | 530 | 0.072 |  | 486 | 0.006 |
|  | 248 | 0.008 |  | 554 | 0.002 |  | 503 | 0.001 |
|  | 390 | 0.008 |  | 556 | 0.001 |  | 506 | 0.365 |
|  | 391 | 0.015 | DUF283 | 598 | 0.001 |  | 507 | 0.001 |
|  | 392 | 0.748 |  | 604 | 0.002 |  | 509 | 0.89 |
|  | 393 | 0.099 |  | 632 | 0.01 |  | 510 | 0.028 |
|  | 394 | 0.27 |  | 633 | 0.001 |  | 511 | 0.029 |
|  | 395 | 0.246 |  | 635 | 0.092 |  | 512 | 0.001 |
|  | 396 | 0.074 |  | 636 | 0.415 |  | 527 | 0.003 |
|  | 397 | 0.047 |  | 639 | 0.508 |  | 529 | 0.022 |
|  | 431 | 0.001 |  | 640 | 0.611 |  | 530 | 0.238 |
|  | 433 | 0.001 |  | 641 | 0.017 |  | 552 | 0.005 |
|  | 434 | 0.003 |  | 642 | 0.111 |  | 556 | 0.004 |
|  | 436 | 0.015 |  | 643 | 0.092 | DUF283 | 632 | 0.001 |
|  | 437 | 0.099 |  | 644 | 0.011 |  | 635 | 0.026 |
|  | 438 | 0.671 |  | 645 | 0.103 |  | 636 | 0.531 |
|  | 439 | 0.012 |  | 647 | 0.001 |  | 639 | 0.559 |
|  | 506 | 0.049 |  | 648 | 0.095 |  | 640 | 0.859 |
|  | 509 | 0.781 |  | 649 | 0.001 |  | 641 | 0.051 |
|  | 510 | 0.052 |  | 712 | 0.031 |  | 642 | 0.195 |
|  | 511 | 0.054 |  | 713 | 0.026 |  | 643 | 0.288 |
|  | 512 | 0.003 |  | 714 | 0.17 |  | 644 | 0.064 |
|  | 525 | 0.001 |  | 715 | 0.026 |  | 645 | 0.216 |
|  | 527 | 0.026 | D-linker | 716 | 0.032 |  | 646 | 0.003 |
|  | 528 | 0.002 |  | 717 | 0.799 |  | 648 | 0.136 |
|  | 529 | 0.018 |  | 718 | 0.328 |  | 649 | 0.002 |
|  | 530 | 0.505 |  | 719 | 0.001 |  | 672 | 0.003 |
|  | 552 | 0.029 |  | 720 | 0.001 |  | 674 | 0.001 |
|  | 554 | 0.022 |  | 725 | 0.016 |  | 707 | 0.002 |
|  | 555 | 0.002 |  | 727 | 0.006 |  | 711 | 0.001 |
|  | 556 | 0.009 |  | 728 | 0.001 |  | 713 | 0.011 |
|  | 557 | 0.001 |  | 729 | 0.019 |  | 714 | 0.212 |
| DUF283 | 632 | 0.001 |  | 732 | 0.014 |  | 715 | 0.186 |
|  | 635 | 0.051 |  | 735 | 0.002 | D-linker | 716 | 0.139 |
|  | 636 | 0.425 |  | 736 | 0.004 |  | 717 | 0.519 |
|  | 638 | 0.001 |  | 737 | 0.227 |  | 718 | 0.188 |
|  | 639 | 0.587 |  | 738 | 0.381 |  | 719 | 0.008 |
|  | 640 | 0.676 |  | 739 | 1.343 |  | 720 | 0.004 |
|  | 641 | 0.024 |  | 740 | 0.029 |  | 721 | 0.002 |
|  | 642 | 0.23 |  | 741 | 0.001 |  | 725 | 0.008 |
|  | 643 | 0.494 |  | 742 | 0.101 |  | 727 | 0.001 |
|  | 644 | 0.122 |  | 743 | 0.488 |  | 738 | 0.021 |
|  | 645 | 0.534 |  | 744 | 0.921 |  | 739 | 1.073 |
|  | 646 | 0.002 |  | 746 | 0.403 |  | 740 | 0.021 |
|  | 647 | 0.017 | Platform | 785 | 0.042 |  | 741 | 0.045 |
|  | 648 | 0.472 |  | 787 | 0.217 |  | 742 | 0.227 |
|  | 649 | 0.015 |  | 788 | 0.244 |  | 743 | 0.444 |
|  | 713 | 0.001 |  | 790 | 0.002 |  | 744 | 0.865 |
|  | 714 | 0.001 |  | 821 | 0.154 |  | 746 | 0.591 |
|  | 715 | 0.001 |  | 859 | 0.092 | Platform | 784 | 0.002 |
| D-linker | 716 | 0.088 |  | 862 | 0.001 |  | 785 | 0.055 |
|  | 717 | 0.869 | PAZ | 934 | 0.001 |  | 787 | 0.034 |
|  | 718 | 0.212 |  | 936 | 0.05 |  | 788 | 0.254 |
|  | 725 | 0.003 |  | 937 | 0.492 |  | 790 | 0.002 |
|  | 730 | 0.013 |  | 938 | 0.071 |  | 821 | 0.087 |
|  | 731 | 0.001 |  | 940 | 0.007 | PAZ | 934 | 0.005 |
|  | 738 | 0.007 |  | 941 | 0.064 |  | 936 | 0.098 |
|  | 739 | 0.893 |  | 943 | 0.315 |  | 937 | 0.338 |
|  | 740 | 0.026 |  | 944 | 0.03 |  | 941 | 0.001 |
|  | 741 | 0.048 |  | 945 | 0.074 |  | 942 | 0.003 |
|  | 742 | 0.263 |  | 960 | 0.241 |  | 943 | 0.167 |
|  | 743 | 0.445 |  | 961 | 0.122 |  | 945 | 0.172 |
|  | 744 | 0.859 |  | 962 | 0.118 |  | 960 | 0.165 |
|  | 746 | 0.514 |  | 964 | 0.023 |  | 961 | 0.065 |
| Platform | 785 | 0.005 |  | 965 | 0.028 |  | 962 | 0.038 |
|  | 787 | 0.254 |  | 968 | 0.033 |  | 964 | 0.018 |
|  | 788 | 0.435 |  | 971 | 0.327 |  | 965 | 0.004 |
|  | 790 | 0.047 |  | 972 | 0.066 |  | 968 | 0.028 |
|  | 821 | 0.047 |  | 975 | 0.446 |  | 971 | 0.327 |
|  | 859 | 0.006 |  | 976 | 0.136 |  | 972 | 0.039 |
| PAZ | 936 | 0.039 |  | 990 | 0.002 |  | 975 | 0.41 |
|  | 937 | 0.418 |  | 991 | 0.007 |  | 976 | 0.127 |
|  | 938 | 0.053 |  | 992 | 0.473 |  | 990 | 0.003 |
|  | 941 | 0.02 |  | 993 | 0.064 |  | 991 | 0.03 |
|  | 942 | 0.001 |  | 994 | 0.042 |  | 992 | 0.6 |
|  | 943 | 0.302 |  | 995 | 0.277 |  | 994 | 0.056 |
|  | 944 | 0.002 |  | 996 | 0.81 |  | 995 | 0.435 |
|  | 945 | 0.079 |  | 997 | 0.001 |  | 996 | 0.733 |
|  | 960 | 0.207 |  | 998 | 0.023 |  | 998 | 0.021 |
|  | 961 | 0.049 |  | 1001 | 0.018 |  | 1001 | 0.05 |
|  | 962 | 0.032 |  | 1002 | 0.004 |  | 1002 | 0.011 |
|  | 964 | 0.007 |  | 1003 | 0.923 |  | 1003 | 0.841 |
|  | 965 | 0.004 |  | 1004 | 0.235 |  | 1004 | 0.023 |
|  | 968 | 0.023 |  | 1005 | 0.007 |  | 1005 | 0.025 |
|  | 971 | 0.28 |  | 1006 | 0.412 |  | 1006 | 0.471 |
|  | 972 | 0.077 |  | 1007 | 0.308 |  | 1007 | 0.574 |
|  | 975 | 0.513 |  | 1008 | 0.328 |  | 1008 | 0.57 |
|  | 976 | 0.117 |  | 1010 | 0.262 |  | 1009 | 0.034 |
|  | 990 | 0.001 |  | 1011 | 0.023 |  | 1010 | 0.411 |
|  | 991 | 0.008 |  | 1012 | 0.849 |  | 1011 | 0.018 |
|  | 992 | 0.357 |  | 1013 | 0.792 |  | 1012 | 0.704 |
|  | 993 | 0.001 |  | 1014 | 0.192 |  | 1013 | 0.542 |
|  | 994 | 0.106 |  | 1015 | 0.592 |  | 1014 | 0.153 |
|  | 995 | 0.236 |  | 1016 | 0.35 |  | 1015 | 0.317 |
|  | 996 | 0.691 |  | 1019 | 0.155 |  | 1016 | 0.051 |
|  | 998 | 0.026 |  | 1020 | 0.327 |  | 1018 | 0.001 |
|  | 1001 | 0.104 |  | 1023 | 0.427 |  | 1019 | 0.414 |
|  | 1002 | 0.016 |  | 1027 | 0.011 |  | 1020 | 0.299 |
|  | 1003 | 0.777 |  | 1028 | 0.161 |  | 1021 | 0.007 |
|  | 1004 | 0.034 |  | 1029 | 0.041 |  | 1022 | 0.024 |
|  | 1005 | 0.02 |  | 1030 | 0.096 |  | 1023 | 0.364 |
|  | 1006 | 0.392 |  | 1031 | 0.681 |  | 1024 | 0.008 |
|  | 1007 | 0.62 |  | 1032 | 0.124 |  | 1027 | 0.006 |
|  | 1008 | 0.57 |  | 1033 | 0.077 |  | 1028 | 0.204 |
|  | 1010 | 0.07 | RIIIDa | 1304 | 0.424 |  | 1029 | 0.204 |
|  | 1011 | 0.013 |  | 1305 | 0.623 |  | 1030 | 0.108 |
|  | 1012 | 0.674 |  | 1307 | 0.275 |  | 1031 | 0.611 |
|  | 1013 | 0.69 |  | 1311 | 0.003 |  | 1032 | 0.154 |
|  | 1014 | 0.23 |  | 1313 | 0.048 |  | 1033 | 0.079 |
|  | 1015 | 0.517 |  | 1314 | 0.001 | RIIIDa | 1303 | 0.003 |
|  | 1016 | 0.333 |  | 1316 | 0.006 |  | 1304 | 0.433 |
|  | 1017 | 0.013 |  | 1324 | 0.013 |  | 1305 | 0.558 |
|  | 1018 | 0.004 |  | 1339 | 0.062 |  | 1307 | 0.26 |
|  | 1019 | 0.372 |  | 1340 | 0.005 |  | 1311 | 0.007 |
|  | 1020 | 0.344 |  | 1341 | 0.055 |  | 1313 | 0.057 |
|  | 1023 | 0.178 |  | 1342 | 0.042 |  | 1314 | 0.001 |
|  | 1024 | 0.002 |  | 1344 | 0.013 |  | 1316 | 0.024 |
|  | 1027 | 0.017 |  | 1345 | 0.35 |  | 1317 | 0.001 |
|  | 1028 | 0.145 |  | 1348 | 0.048 |  | 1324 | 0.023 |
|  | 1029 | 0.106 |  | 1349 | 0.022 |  | 1339 | 0.189 |
|  | 1030 | 0.109 |  | 1351 | 0.003 |  | 1341 | 0.23 |
|  | 1031 | 0.787 |  | 1352 | 0.014 |  | 1342 | 0.183 |
|  | 1032 | 0.123 |  | 1353 | 0.021 |  | 1344 | 0.017 |
|  | 1033 | 0.074 |  | 1354 | 0.002 |  | 1345 | 0.47 |
| RIIIDa | 1303 | 0.007 |  | 1358 | 0.001 |  | 1347 | 0.005 |
|  | 1304 | 0.5 |  | 1371 | 0.001 |  | 1348 | 0.315 |
|  | 1305 | 0.551 |  | 1372 | 0.001 |  | 1349 | 0.072 |
|  | 1307 | 0.318 |  | 1373 | 0.502 |  | 1351 | 0.006 |
|  | 1309 | 0.03 |  | 1374 | 0.001 |  | 1352 | 0.022 |
|  | 1310 | 0.001 |  | 1375 | 0.107 |  | 1353 | 0.23 |
|  | 1311 | 0.031 |  | 1378 | 0.004 |  | 1354 | 0.005 |
|  | 1313 | 0.124 |  | 1379 | 0.001 |  | 1373 | 0.442 |
|  | 1314 | 0.088 |  | 1541 | 0.117 |  | 1374 | 0.001 |
|  | 1316 | 0.037 |  | 1542 | 0.231 |  | 1375 | 0.012 |
|  | 1317 | 0.001 |  | 1543 | 0.042 |  | 1541 | 0.088 |
|  | 1320 | 0.001 |  | 1544 | 0.008 |  | 1542 | 0.34 |
|  | 1324 | 0.05 |  | 1546 | 0.006 |  | 1543 | 0.036 |
|  | 1339 | 0.044 |  | 1555 | 0.002 |  | 1544 | 0.001 |
|  | 1340 | 0.002 |  | 1557 | 0.105 |  | 1555 | 0.015 |
|  | 1341 | 0.046 |  | 1558 | 0.004 |  | 1557 | 0.555 |
|  | 1342 | 0.038 |  | 1561 | 0.004 |  | 1558 | 0.004 |
|  | 1344 | 0.014 | RIIIDb | 1702 | 0.001 |  | 1561 | 0.026 |
|  | 1345 | 0.249 |  | 1728 | 0.002 | RIIIDb | 1702 | 0.001 |
|  | 1347 | 0.012 |  | 1729 | 0.017 |  | 1728 | 0.004 |
|  | 1348 | 0.313 |  | 1730 | 0.132 |  | 1729 | 0.091 |
|  | 1349 | 0.06 |  | 1731 | 0.008 |  | 1730 | 0.488 |
|  | 1351 | 0.013 |  | 1733 | 0.033 |  | 1731 | 0.025 |
|  | 1352 | 0.021 |  | 1734 | 0.107 |  | 1733 | 0.346 |
|  | 1353 | 0.252 |  | 1736 | 0.003 |  | 1734 | 0.231 |
|  | 1354 | 0.001 |  | 1737 | 0.03 |  | 1737 | 0.301 |
|  | 1373 | 0.54 |  | 1740 | 0.001 |  | 1740 | 0.017 |
|  | 1374 | 0.001 |  | 1741 | 0.003 |  | 1741 | 0.035 |
|  | 1375 | 0.037 |  | 1785 | 0.111 |  | 1742 | 0.002 |
|  | 1541 | 0.051 |  | 1806 | 0.009 |  | 1785 | 0.072 |
|  | 1542 | 0.107 | dsRBD | 1851 | 0.046 |  | 1786 | 0.001 |
|  | 1543 | 0.026 |  | 1852 | 0.358 |  | 1806 | 0.094 |
|  | 1544 | 0.007 |  | 1853 | 0.004 |  | 1849 | 0.001 |
|  | 1546 | 0.008 |  | 1854 | 0.237 | dsRBD | 1851 | 0.062 |
|  | 1555 | 0.005 |  | 1855 | 1.42 |  | 1852 | 0.33 |
|  | 1557 | 0.516 |  | 1858 | 0.363 |  | 1853 | 0.003 |
|  | 1558 | 0.001 |  | 1859 | 0.043 |  | 1854 | 0.243 |
|  | 1561 | 0.022 |  | 1862 | 0.01 |  | 1855 | 1.454 |
| RIIIDb | 1728 | 0.012 |  | 1863 | 0.004 |  | 1858 | 0.337 |
|  | 1729 | 0.158 |  | 1866 | 0.008 |  | 1859 | 0.041 |
|  | 1730 | 0.478 |  | 1867 | 0.011 |  | 1862 | 0.031 |
|  | 1731 | 0.033 |  | 1869 | 0.004 |  | 1863 | 0.032 |
|  | 1732 | 0.001 |  | 1870 | 0.141 |  | 1865 | 0.001 |
|  | 1733 | 0.342 |  | 1871 | 0.001 |  | 1866 | 0.027 |
|  | 1734 | 0.213 |  | 1872 | 1.086 |  | 1867 | 0.111 |
|  | 1736 | 0.029 |  | 1873 | 0.02 |  | 1869 | 0.008 |
|  | 1737 | 0.314 |  | 1874 | 0.047 |  | 1870 | 0.323 |
|  | 1740 | 0.011 |  | 1875 | 0.12 |  | 1871 | 0.001 |
|  | 1741 | 0.02 |  | 1876 | 0.179 |  | 1872 | 1.376 |
|  | 1742 | 0.001 |  | 1877 | 0.005 |  | 1873 | 0.007 |
|  | 1785 | 0.242 |  | 1878 | 0.097 |  | 1874 | 0.095 |
|  | 1806 | 0.131 |  | 1895 | 0.527 |  | 1875 | 0.004 |
| dsRBD | 1851 | 0.091 |  | 1896 | 0.3 |  | 1876 | 0.092 |
|  | 1852 | 0.324 |  | 1897 | 0.305 |  | 1877 | 0.001 |
|  | 1854 | 0.236 |  | 1898 | 1.142 |  | 1878 | 0.127 |
|  | 1855 | 1.519 |  | 1899 | 0.008 |  | 1895 | 0.662 |
|  | 1858 | 0.363 |  | 1901 | 0.353 |  | 1896 | 0.276 |
|  | 1859 | 0.077 |  | 1902 | 0.05 |  | 1897 | 0.356 |
|  | 1862 | 0.011 |  |  |  |  | 1898 | 1.281 |
|  | 1863 | 0.005 |  |  |  |  | 1899 | 0.011 |
|  | 1865 | 0.001 |  |  |  |  | 1901 | 0.364 |
|  | 1866 | 0.012 |  |  |  |  | 1902 | 0.049 |
|  | 1867 | 0.023 |  |  |  |  |  |  |
|  | 1869 | 0.002 |  |  |  |  |  |  |
|  | 1870 | 0.035 |  |  |  |  |  |  |
|  | 1872 | 0.926 |  |  |  |  |  |  |
|  | 1873 | 0.002 |  |  |  |  |  |  |
|  | 1874 | 0.004 |  |  |  |  |  |  |
|  | 1876 | 0.037 |  |  |  |  |  |  |
|  | 1878 | 0.047 |  |  |  |  |  |  |
|  | 1895 | 0.359 |  |  |  |  |  |  |
|  | 1896 | 0.305 |  |  |  |  |  |  |
|  | 1897 | 0.571 |  |  |  |  |  |  |
|  | 1898 | 1.459 |  |  |  |  |  |  |
|  | 1899 | 0.007 |  |  |  |  |  |  |
|  | 1901 | 0.371 |  |  |  |  |  |  |
|  | 1902 | 0.067 |  |  |  |  |  |  |
